# miR-184 modulates *dilp8* to control developmental timing during normal growth conditions and in response to developmental perturbations

**DOI:** 10.1101/2024.07.08.602443

**Authors:** Jervis Fernandes, Muhammed Naseem, Ayisha Marwa MP, Jishy Varghese

## Abstract

Organismal development depends on the precise coordination of growth and developmental timing, which is regulated by a complex interplay of intrinsic and extrinsic factors. However, the mechanisms underlying this regulation are not fully understood. Post-transcriptional regulation by microRNAs (miRNAs) plays a pivotal role in ensuring the proper timing of gene expression necessary for growth and development. In this study, we conducted a genetic screen to identify microRNAs that regulate developmental timing in *Drosophila*. Our screen identified *miR-184,* previously implicated in germline maturation and embryonic development, as a regulator of pupariation timing by acting in the larval imaginal discs. Using genetic and molecular approaches, we identified *Drosophila insulin-like peptide 8* (*Dilp8*), a paracrine factor critical for regulating developmental stability, as a target of *miR-184*. During normal larval development miR-184 facilitates timely pupariation by regulating *dilp8* levels. Furthermore, we demonstrate that miR-184 plays a critical role in tissue damage responses by inducing *dilp8* expression, which delays pupariation to enable damage repair mechanisms. These findings reveal a novel post-transcriptional regulatory mechanism that links miR-184 to the control of developmental timing under normal growth conditions and in response to tissue damage.

## Introduction

Normal growth and development are essential for an organism’s proper functioning and reproductive capacity. Disruptions in these processes can lead to anomalies such as developmental defects, metabolic imbalances, and diseases like cancer. Environmental factors, including nutrition and stress, significantly influence growth and development. In *Drosophila*, growth and developmental timing are tightly linked, with reaching a critical body weight being necessary for the transition from larval to pupal stages, which triggers adult metamorphosis. Understanding how growth and developmental timing are regulated in *Drosophila* can provide insights into similar mechanisms in other animals, including humans, and suggest potential therapeutic interventions for related developmental disorders.

In *Drosophila*, transitions between larval instars and from larva to pupa are initiated by pulses of the molting hormone ecdysone (Baehrecke, 1996; Yamanaka et al., 2013). Ecdysone is secreted by the prothoracic gland and converted into its active form, 20-hydroxyecdysone (20-HE), in the larval fat body and other tissues. This hormone initiates a transcriptional cascade via the EcR/USP receptor complex, triggering metamorphosis and halting growth (Yamanaka et al., 2013). The timing of pupariation is governed by ecdysone secretion; reduced or delayed ecdysone pulses disrupt development and delay pupariation (Baehrecke, 1996; Yamanaka et al., 2013). Such delays extend the larval feeding period, leading to excessive growth. Environmental factors like temperature, crowding, nutrition, oxygen availability, and toxins can modulate pupariation timing (Beamish et al., 2021; Bonnier, 1926; Klepsatel et al., 2018; Layalle et al., 2008; Turingan et al., 2024). Insulin signaling in the prothoracic gland plays a crucial role in ecdysone production and metamorphosis by modulating the expression of the microRNA Bantam (Boulan et al., 2013). Additionally, 20-HE and juvenile hormone (JH) exhibit antagonistic relationships, with JH inhibiting ecdysteroid biosynthesis and being inhibited by 20-HE (Liu et al., 2018). Nutrient stress also modulates ecdysone secretion via AKH (Adipokinetic hormone, *Drosophila* equivalent of Glucagon), which regulates calcium signaling in the prothoracic gland (Hughson et al., 2021). Thus, pupariation timing is controlled by both internal and external factors acting on the prothoracic gland.

A bilateral pair of neurons in the larval central brain release prothoracicotropic hormone (PTTH), which stimulates the prothoracic gland to produce ecdysone (McBrayer et al., 2007; Ghosh et al., 2010). Loss of the *ptth* gene leads to delayed pupariation and increased body size due to prolonged feeding. PTTH release is regulated by upstream neuronal inputs that control the timing of PTTH secretion, affecting developmental timing and the onset of metamorphosis (Ghosh et al., 2010; McBrayer et al., 2007). Ecdysteroid biosynthesis involves cytochrome P450 monoxygenases encoded by Halloween genes, which are tightly regulated during ecdysone synthesis. The transcript levels of Halloween genes correlate well with the ecdysone titer, implying a stringent transcriptional regulation of these genes during ecdysone synthesis. Disruptions in Halloween gene expression lead to delayed pupariation (Kamiyama & Niwa, 2022). Additionally, transcriptional regulators that act across various signaling pathways modulate Halloween genes (Danielsen et al., 2014; Kamiyama & Niwa, 2022).

Recent studies also highlight the role of imaginal discs in regulating pupariation timing. Imaginal discs secrete factors like DILP8 (*Drosophila* insulin-like peptide-8), Dpp (Decapentaplegic), and Upd3 (Unpaired-3) to influence pupariation timing. DILP8, a relaxin-like peptide, buffers developmental noise and delays pupariation in response to growth defects in the imaginal discs (Colombani et al., 2012; Katsuyama et al., 2015). Growth perturbations in the imaginal discs trigger JNK pathway activity, upregulating DILP8 levels in the hemolymph (Colombani, Andersen, & Leopold, 2012; Katsuyama et al., 2015). DILP8 delays pupariation by acting on Lgr3 receptors in the larval brain, which connect PTTH neurons to the prothoracic gland, regulating ecdysone signaling (Colombani, Andersen, & Leopol, 2012, 2012; Garelli et al., 2015; Katsuyama et al., 2015; Vallejo et al., 2015). Furthermore, damage to the larval hindgut increases DILP8 levels, contributing to delayed pupariation (Cohen et al., 2021). Dpp inhibits ecdysone biosynthesis to prevent premature metamorphosis during imaginal disc growth, while reduced Dpp levels allow progression to metamorphosis once disc growth is complete (Setiawan et al., 2018). Upd3, a cytokine secreted by the imaginal discs, delays pupariation in response to malignant transformations by upregulating Bantam and activating JAK/STAT signaling (Romao et al., 2021).

MicroRNAs also play a crucial role in regulating metamorphosis. For example, bantam microRNA in the prothoracic gland modulates pupariation timing by influencing ecdysone levels (Boulan et al., 2013). Bantam also controls the expression of *jhamt*, a gene involved in JH biosynthesis in the corpus allatum (Qu et al., 2017). miR-8, expressed in the corpus allatum, regulates JH biosynthesis and pupariation timing (Zhang et al., 2021). Numerous microRNAs are differentially modulated by ecdysone during the larval-pupal transition, highlighting their roles in the regulation of metamorphosis (Jin et al., 2020; Lim et al., 2018). However, the role of microRNAs in regulating the onset of metamorphosis in response to tissue damage has yet to be fully explored.

We report that miR-184, a conserved microRNA critical for oogenesis and early embryonic development in *Drosophila* (Iovino et al., 2009), plays a significant role in regulating developmental timing. *miR-184* mutants exhibit delayed pupariation due to reduced ecdysone signaling. We show that *miR-184* functions in healthy imaginal discs to regulate pupariation timing by targeting *dilp8*. Our results also reveal that modulation of *miR-184* in response to imaginal disc damage delays the larval-to-pupal transition by upregulating *dilp8* levels. These findings underscore the essential role of miR-184 in controlling developmental timing both under normal growth conditions and in response to tissue damage.

## Results

### Role of miR-184 in the regulation of pupariation timing

MicroRNAs (miRNAs) are known to regulate various developmental processes, including the timing of developmental transitions. In this study, we explored the role of *Drosophila* miRNAs in regulating the timing of pupariation, a critical stage when the larva enters the metamorphosis stage under the influence of ecdysone hormone (Table S1). While several miRNAs caused delays in pupariation by 12 hours or more, we focused on miR-184, as it is the highest-expressed miRNA in the larvae (FlyAtlas 2.0) and the ring gland (Zhang et al., 2021). We observed that homozygous mutants of *miR-184* exhibited significant delays in puparium formation, indicating that *miR-184* is essential for the proper timing of the larval-to-pupal transition (Fig. 1A). To further elucidate the function of miR-184, we employed *miR-184-sponge (miR-184-sp)*, a transgenic inhibitor of *miR-184* (Fulga et al., 2015), to reduce *miR-184* expression in the entire organism using the *tubGAL4* driver line. Similar to the mutants, downregulation of *miR-184* in the entire organism resulted in a delay in pupariation (Fig. S1; Fig. 1B). The delay in pupariation was rescued by co-overexpression of *miR-184*, confirming the specificity of the *miR-184 sponge* (Fig. S2). These results revealed that miR-184 regulates the timing of pupariation, which prompted us to investigate the regulation of pupariation timing by miR-184.

**Figure 1.**
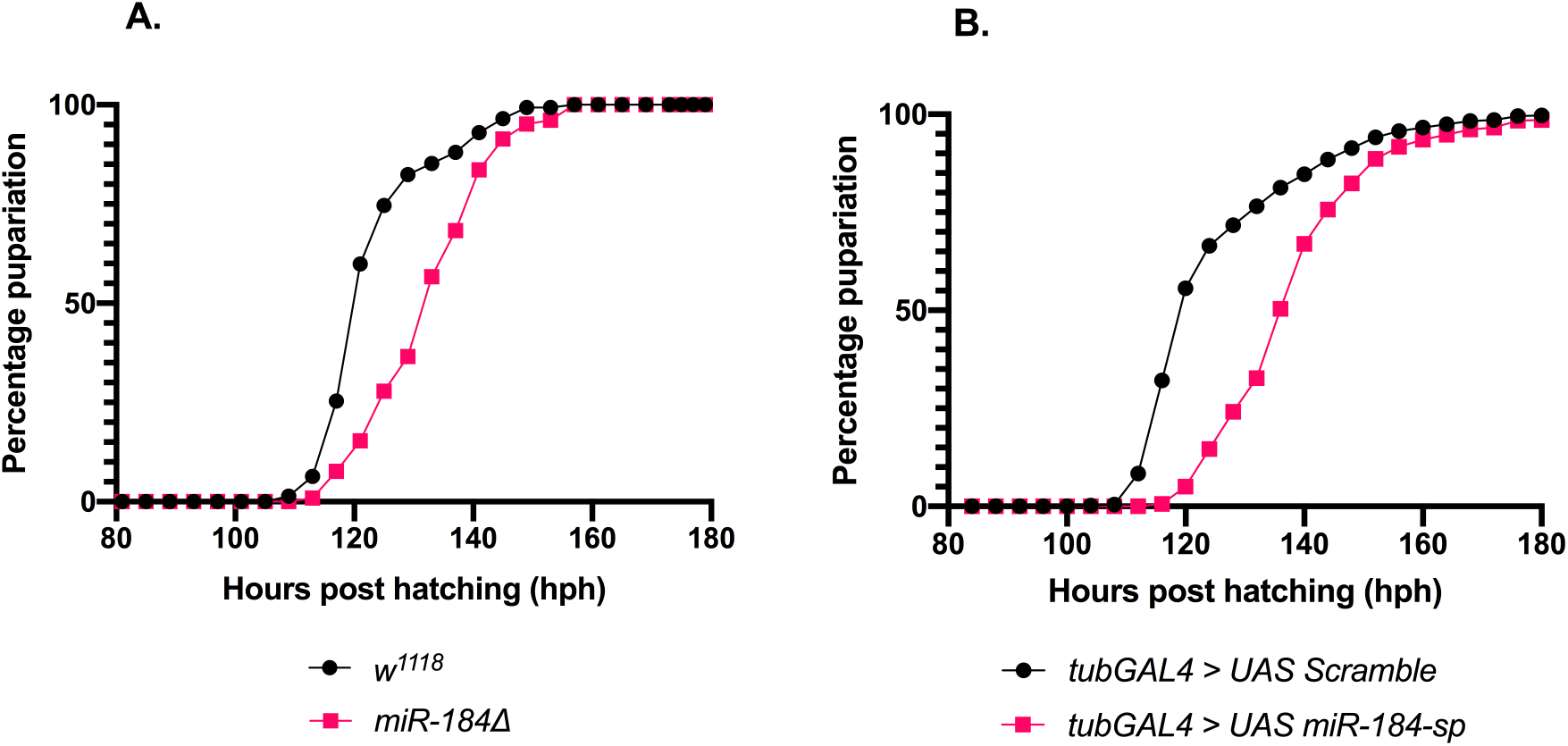
miR-184 plays a role in the regulation of pupariation time point. (A) *miR-184* mutants display delayed pupariation, pupariation rates of *w^1118^* (control) (n=142) and *miR-184* homozygous null mutants (n=104). Data from three independent experiments are shown; (p<0.0001). (B) Downregulation of *miR-184* leads to delayed pupariation, pupariation of *tubGal4, UAS Scramble* (control) (n=955), and *tubGal4, UAS-miR-184-sp* (n*=*617). Data from three independent experiments are shown for A and B; (p<0.0001). Data points plotted are the percentage of larvae pupariated, LogRank test was performed to determine statistical significance.

### miR-184 manages optimal Ecdysone signaling

The timing of pupariation is tightly controlled by the steroid hormone ecdysone, which is released by the prothoracic gland in developing larvae. Ecdysone is converted into its active form, 20-hydroxyecdysone (20-HE), in peripheral tissues, and regulates the expression of ecdysone-responsive genes, critical for pupariation, metamorphosis, and adult emergence. Given the pupariation delay observed in *miR-184*-depleted larvae, we hypothesized that miR-184 might be involved in maintaining optimal ecdysone signaling.

To test this, we first measured the expression of genes that promote ecdysone biosynthesis: *prothoracicotropic hormone* (*ptth*), *phantom* (*phm*), and *disembodied* (*dib*) in response to downregulation of *miR-184* in the whole larva. We found that *miR-184* downregulation led to a significant decrease in the mRNA levels of these genes (Fig. 2A). Furthermore, the transcript levels of key ecdysone-response genes- *Ecdysone inducible protein 74* (*E74*), *Ecdysone inducible protein 75* (*E75*), *Broad-complex* (*BR-C*), and *fatbody protein-1* (*fbp1*), was also reduced in *miR-184*-depleted larvae (Fig. 2B). We then proceeded to measure the levels of 20-hydroxyecdysone (20-HE), in the late 3rd instar larvae. Furthermore, the level of 20-HE was found to be significantly lower in *miR-184*-depleted larvae (Fig. 2C). To confirm the role of reduced ecdysone signaling in the pupariation delay, we fed 20-HE to *miR-184*-depleted larvae, which completely rescued the pupariation delay (Fig. 2D). These results confirm that miR-184 is required for optimal ecdysone signaling and the timely transition from larva to pupa. Next, we focused on identifying the tissue where *miR-184* acts in regulating ecdysone signaling.

**Figure 2.**
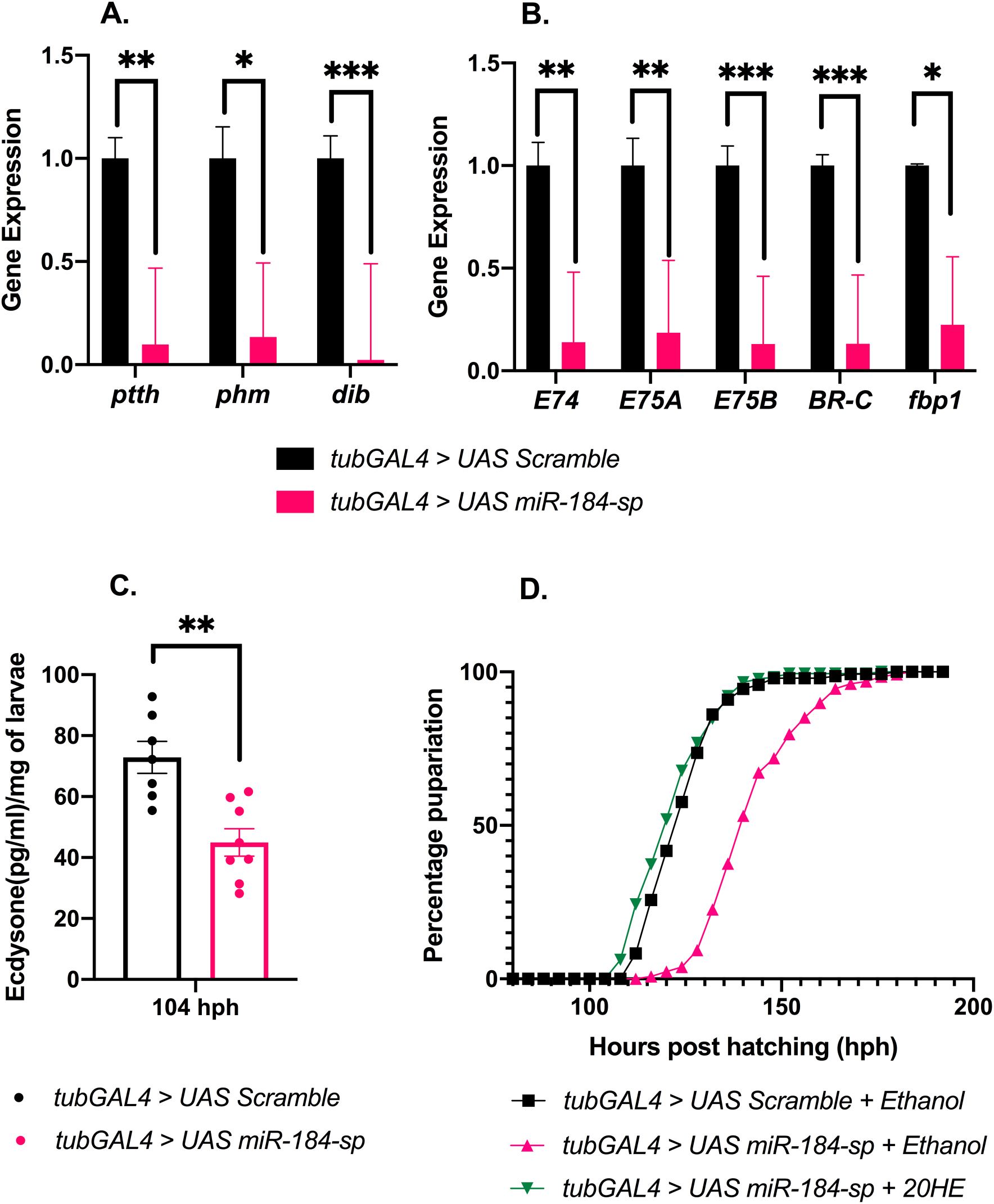
miR-184 levels maintain proper Ecdysone signaling. (A) Expression of Ecdysone biosynthesis genes in response to ubiquitous downregulation of *miR-184*; using *tubGal4, UAS-miR-184-sp* and *tubGal4, UAS Scramble* (control), from three independent experiments are shown. Fold change of transcript levels of *ptth*, *phm,* and *dib* are shown, normalized to *rp49*. Total RNA was extracted from whole larvae. Bars graphs plotted are average/mean values, error bars represent SEM; a two-way ANOVA was performed to determine statistical significance; * p < 0.05, ***p<0.001, ****p<0.0001. (B) Expression of Ecdysone response genes in response to ubiquitous downregulation of *miR-184* using *tubGal4*; *tubGal4, UAS-Scramble* (control) and *tubGal4, UAS-miR-184-sponge*, data from three independent experiments are shown. Fold change of transcript levels of *E74*, *E75A*, *E75B*, *BR-C,* and *fbp1* are shown, normalized to *rp49*. Total RNA was extracted from whole larvae. Bars graphs plotted are average/mean values, error bars represent SEM; a two-way ANOVA was performed to determine statistical significance; * p < 0.05, ***p<0.001, ****p<0.0001. (C) Measurement of 20-Hydroxyecdysone levels (pg/ml)/mg of the larvae at 104 hours post-hatching in *tubGal4, UAS-Scramble* (control) and *tubGal4, UAS-miR-184-sponge*, data from seven independent experiments are shown. Bars graphs plotted are absolute values, error bars represent SEM; a Welch’s t-test was performed to determine statistical significance; * p < 0.05, ***p<0.001, ****p<0.0001. (D) Pupariation time of *tubGAL4> miR-184-sp* larvae fed 20-Hydroxyecdysone (0.3 mg/mL) (p=0.0748) or ethanol (Ethanol) (p<0.0001) in comparison to *tubGAL4> control-sp* larvae (*n* ≥128 larvae per treatment). Log rank test was carried out to determine significance.

### miR-184 acts in the imaginal discs to regulate pupariation timing

Although *miR-184* is highly expressed in the ring gland (Zhang et al., 2021), we observed no pupariation delay when *miR-184* was downregulated in the prothoracic gland, the primary source of ecdysone (Fig. 3A). This indicates that the pupariation delay in *miR-184* mutants is not due to *miR-184* depletion in the prothoracic gland. Similarly, downregulation of *miR-184* in the corpus allatum (CA), which produces juvenile hormone (JH) and negatively regulates ecdysone, did not affect pupariation timing (Fig. 3B). Moreover, downregulation of *miR-184* in the PTTH neurons in the brain, which regulate ecdysone production, also failed to alter pupariation timing (Fig. 3C). These findings suggest that the role of *miR-184* in regulating pupariation timing is not mediated through these tissues. To identify the site of *miR-184* activity, we focused on other peripheral tissues where *miR-184* could act and regulate ecdysone signaling.

**Figure 3.**
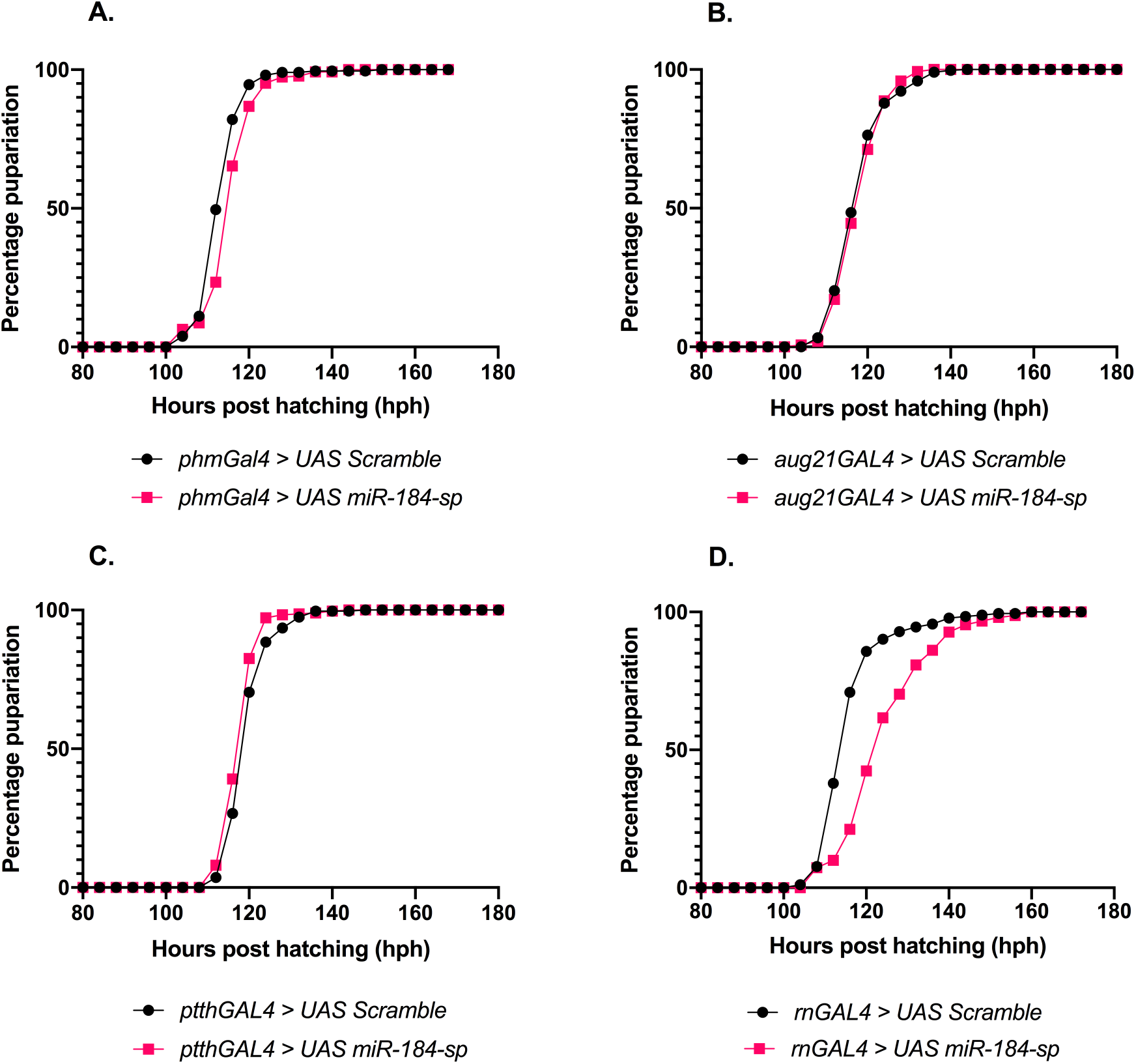
miR-184 levels in the imaginal discs regulate pupariation timepoint. (A) Downregulation of *miR-184* in the prothoracic gland leads to minimal effects on pupariation time point. *phmGal4, UAS-miR-184-sp*, (n*=*206) and *phmGal4, UAS Scramble* (control) (n*=*265), three independent experiments are shown; p<0.0001. (B) Downregulation of *miR-184* in the corpus allatum leads to minimal effects on pupariation time point. *aug21Gal4, UAS-miR-184-sp* (n=601) and *aug21Gal4, UAS Scramble* (control) (n*=*292), six independent experiments are shown; p<0.0001. (C) Downregulation of miR-184 in the PTTH neurons leads to minimal effects on pupariation time point. *ptthGal4, UAS Scramble* (control) (n*=*277) and *ptthGal4, UAS-miR-184-sp (n=286)*, three independent experiments are shown; p<0.0001. (D) Downregulation of *miR-184* in the imaginal discs leads to a delay in pupariation timepoint. *rotundGal4, UAS-miR-184-sponge* (n=151), and *rotundGal4, UAS-Scramble* (control) (n*=*182), three independent experiments are shown; p<0.0001. Data points plotted are the percentage of larvae pupariated, LogRank test was performed to determine statistical significance (A-D).

Recent studies report that adult imaginal discs in larvae play a role in regulating the timing of pupariation, especially in response to tissue perturbations. When *miR-184* was downregulated in the imaginal discs using the *rnGAL4* driver line, we observed a significant delay in pupariation (Fig. 3D). Furthermore, *miR-184* depletion in the imaginal discs reduced the expression of ecdysone biosynthesis and ecdysone-response genes (Fig. S3A, B). These results indicate that miR-184 functions in the imaginal discs to regulate ecdysone signaling and the timing of pupariation. As microRNAs function by regulating target gene expression, next we sought to identify *miR-184* target genes in the imaginal discs that influence pupariation timing.

### miR-184 modulates pupariation timing via its target *dilp8*

Our target prediction analysis identified *dilp8* as a potential target of *miR-184* (Fig. S4). The imaginal discs have been shown to regulate pupariation timing, especially in response to growth perturbations (Colombani, Andersen, & Leopold, 2012; Garelli et al., 2012; Romao et al., 2021; Setiawan et al., 2018). One of the key factors involved in this regulation is DILP8, which acts as a developmental checkpoint to delay pupariation in response to growth perturbations (Colombani et al., 2012; Garelli et al., 2012). We hypothesized that *miR-184* regulates pupariation timing by modulating *dilp8* expression.

We found that *dilp8* mRNA levels were elevated in both *miR-184* knockout larvae (Fig. 4A) and in larvae with whole-body downregulation of *miR-184* (Fig. 4B), indicating that *dilp8* could be a target of *miR-184*. This suggests that *miR-184* restricts *dilp8* expression during larval development and regulates pupariation timing. To confirm this, we tested whether miR-184 directly regulates *dilp8* via its 3’ untranslated region (UTR) by using a *dilp8-3’UTR-GFP* reporter. We observed reduced GFP expression in the wing imaginal discs in response to *miR-184* overexpression, confirming that miR-184 can negatively regulate *dilp8* via its 3’UTR (Fig. 4C-D’).

**Figure 4.**
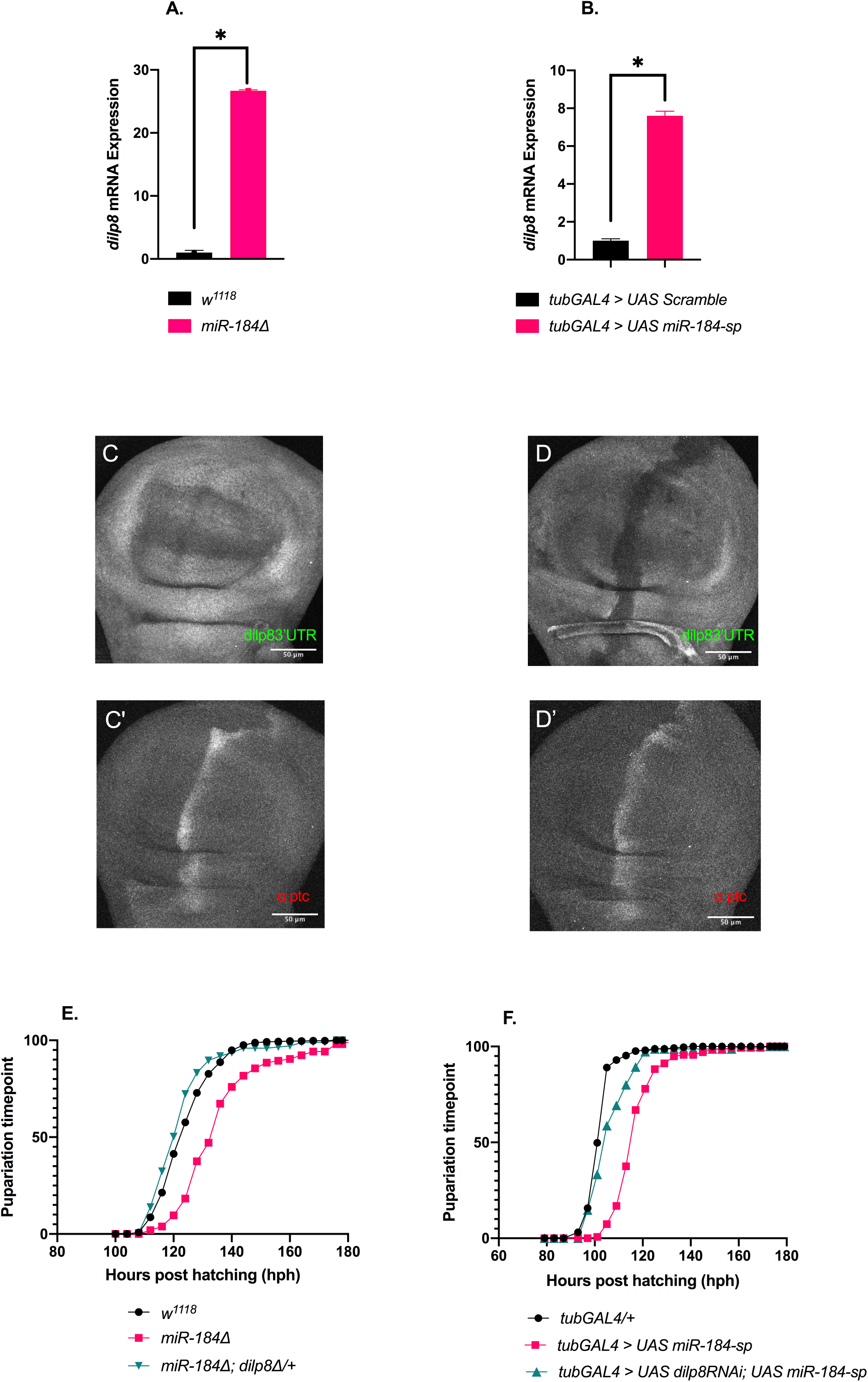
miR-184 regulates developmental time point via its target gene dilp8. (A) *dilp8* mRNA levels in *w^1118^* (control), and *miR-184* homozygous null mutants from four independent experiments are shown; p=0.0281. Total RNA was extracted from whole larvae. (B) *dilp8* mRNA levels in *tubGal4, UAS Scramble* (control),and *tubGal4, UAS-miR-184-sp*, four independent experiments are shown; p=0.0428. Total RNA was extracted from whole larvae. Bars graphs plotted are average/mean values, error bars represent SEM, Welch’s t-test was performed to determine statistical significance (for A and B) (C) *dilp8* 3’UTR-GFP reporter assay, reporter GFP expression (anti-GFP, green) in control (*ptcGal4/+)* and (D) reporter GFP expression (anti-GFP, green) in *ptcGal4, UAS-miR-184 OE.* (C’) Patched expression (anti-Ptc, red) in *ptcGal4* (control) and (D’) Patched expression (anti-Ptc, red) in *ptcGal4, UAS-miR-184* (overexpression). (E) Rescue of pupariation delay, *w1118* (control) (n=486); *miR-184^-/-^* (homozygous mutant) (n=104) and *miR-184^-/-^; dilp8^-/+^* (rescue) (n=173), from three independent experiments are shown. (F) Rescue of pupariation delay, *tubGal4, UAS Scramble* (control) (n=255) and *tubGal4, UAS-miR-184-sp* (*miR-184* downregulation) (n=136) and *tubGal4, UAS-miR-184-sp, dilp8-RNAi* (rescue) (n=76), three independent experiments are shown. Data points plotted are percentage pupariated, LogRank test was performed (E and F) to determine statistical significance. * p < 0.05, ***p<0.001, ****p<0.0001.

We then tested whether elevated *dilp8* levels in *miR-184* mutants were responsible for the pupariation delay. Indeed, reducing *dilp8* levels in *miR-184* mutant larvae by removing one copy of *dilp8* rescued the pupariation delay (Fig. 4E). Similarly, coexpression of *dilp8 RNAi* with *miR-184 sponge* also rescued the pupariation timing (Fig. 4F). These results confirm that *miR-184* regulates pupariation timing by controlling *dilp8* expression during normal development.

### miR-184 regulates a developmental checkpoint in response to perturbations

In response to developmental stresses, such as excessive cell death or hyperplastic growth, *dilp8* expression is upregulated, which suppresses ecdysone synthesis, delaying pupariation to allow for tissue repair and growth coordination (Katsuyama et al., 2015; Garelli et al., 2015; Vallejo et al., 2015). To investigate whether miR-184 contributes to developmental stress responses by aiding in DILP8 induction, we induced tissue damage in the imaginal discs using a cold-sensitive version of the type 2 ribosome-inactivating protein Ricin-A (RA^CS^), which causes cell death and elevates *dilp8* expression (Sanchez et al., 2019). *Ricin-A^CS^* expression in the imaginal discs led to increased *dilp8* levels and a significant delay in pupariation (Fig. 5A, B). We also observed that *RA^CS^* expression activated JNK signaling, a known pathway involved in the stress response and elevation of *dilp8* expression (Sanchez et al., 2019), by assessing the expression of *puc* (*puckered*) and *mmp1* (*matrix metalloproteinase 1*) - two target genes of JNK signaling - in the wing imaginal discs (Fig. 5C, D).

**Figure 5.**
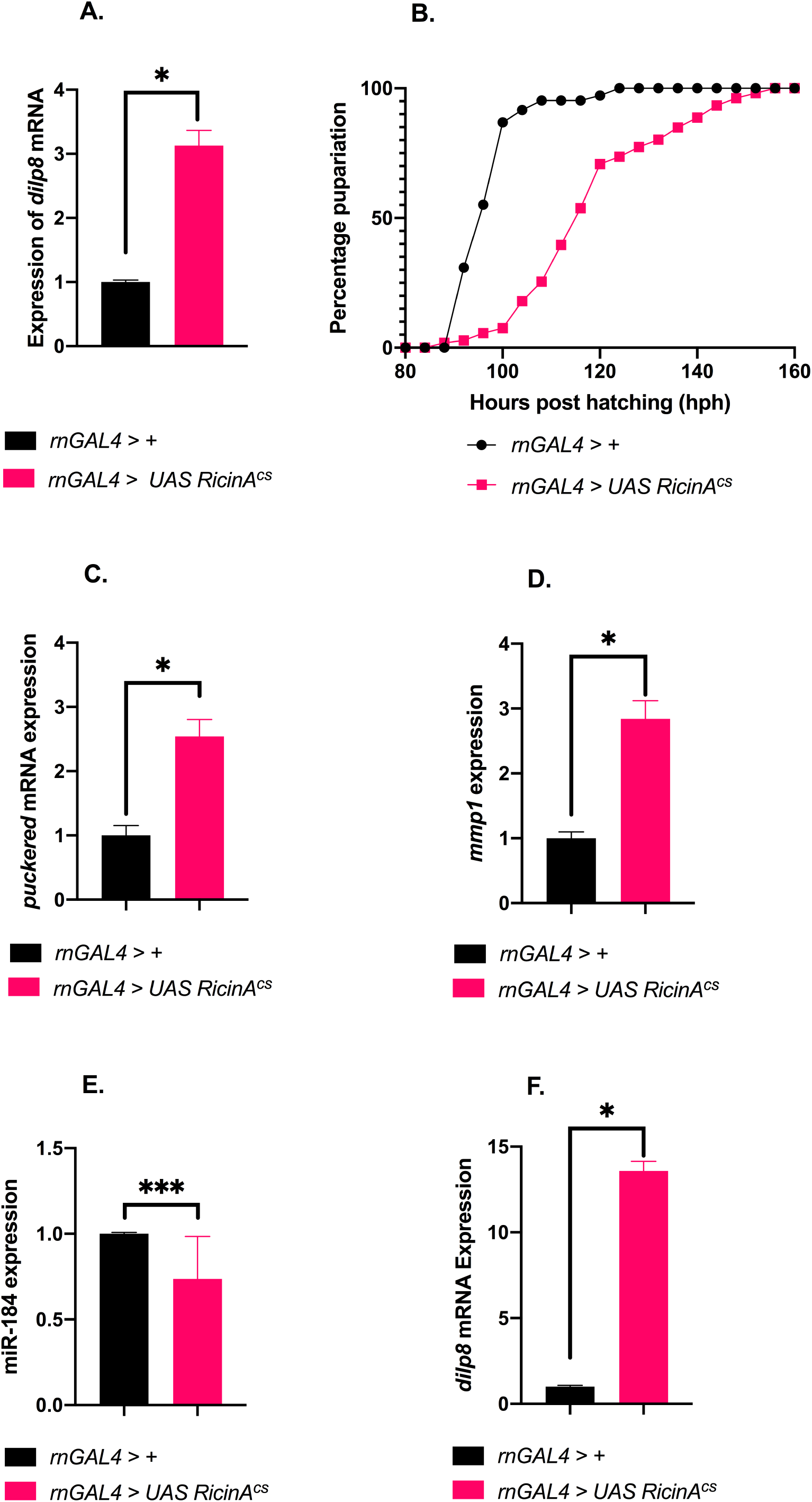
Effects of Ricin-induced developmental perturbation. (A) *dilp8* mRNA levels increased when Ricin-A was expressed in the imaginal discs, *rnGal4* (control) and *rnGal4, UAS-RA^cs^,* three independent experiments are shown; p<0.0243. Fold change of transcript levels of *dilp8* is shown, normalized to *rp49*. Total RNA was extracted from the whole larvae. Bars graphs plotted are average/mean values, error bars represent SEM, and a Welch’s t-test was carried out to determine statistical significance (B) Pupariation delays were observed when Ricin-A*^cs^*was expressed in the imaginal discs, pupariation rates of *rnGal4* (control), n=107 and *rnGal4, UAS-RA^cs^,* n*=*159, three independent experiments are shown; p<0.0001. Data points are percentage pupariated, LogRank test was performed to determine statistical significance. (C-D) JNK target levels increase when Ricin-A was expressed in the imaginal discs, (C) *puckered* mRNA levels in *rnGal4* (control) and *rnGal4, UAS-RA^cs^*, four independent experiments are shown; p= 0.0383. (D) *mmp1* mRNA levels in *rnGal4* (control), and *rnGal4, UAS-RA^cs^*, three independent experiments are shown; p=0.0172. Fold change of transcript levels of *puc* and *mmp1* are shown, normalized to *rp49*. (E) miR-184 levels are affected in response to Ricin-A expression in the imaginal discs, *rnGal4/+* (control) and *rnGal4, UAS-RA^cs^*, four independent experiments are shown; p=0.001. Fold change of transcript levels of *miR-184* is shown, normalized to *2SrRNA*. (F) *dilp8* mRNA levels increase when Ricin-A was expressed in the imaginal discs, *rnGal4/+* (control) and *rnGal4, UAS-RA^cs^,* three independent experiments are shown; p=0.0417. Fold change of transcript levels of *dilp8* is shown, normalized to *rp49*, For C-F, Total RNA was extracted from the wing imaginal discs. Bars graphs plotted are average/mean values, error bars represent SEM, and a Welch’s t-test was performed to determine statistical significance.* p < 0.05, ***p<0.001, ****p<0.0001.

Despite the well-documented dynamic expression pattern of *miR-184* during larval development and its regulation by nutrient availability and tumorous conditions (Fernandes et al., 2022; Shu et al., 2017), its modulation during developmental perturbations remains unexplored. Our findings imply that *miR-184* plays a role in inducing *dilp8* expression in response to such perturbations. We tested this by first measuring the levels of *miR-184* in response to tissue damage induced by *RA^CS^* expression. Remarkably, *RA^CS^* expression in imaginal discs led to a decrease in *miR-184* levels, which corresponded with increased *dilp8* expression (Fig. 5E, F). This finding suggests that *miR-184* plays a role in the responses to developmental stress.

To confirm that reduced *miR-184* expression contributes to *dilp8* upregulation and the pupariation delay under stress, we manipulated *dilp8* expression in *RA^CS^* -induced conditions. Downregulating *dilp8* using RNAi in the *RA^CS^* overexpression background reduced *dilp8* expression (Fig. 6A) and rescued the pupariation delay (Fig. 6C). Moreover, co-overexpression of *miR-184* with *RA^CS^* suppressed *dilp8* expression (Fig. 6B) and rescued the pupariation delay (Fig. 6C). These experiments confirm that *miR-184* modulates *dilp8* expression to mediate pupariation timing under conditions of tissue damage. Thus, our results show that developmental tissue damage responses act through the regulation of *miR-184*.

**Figure 6.**
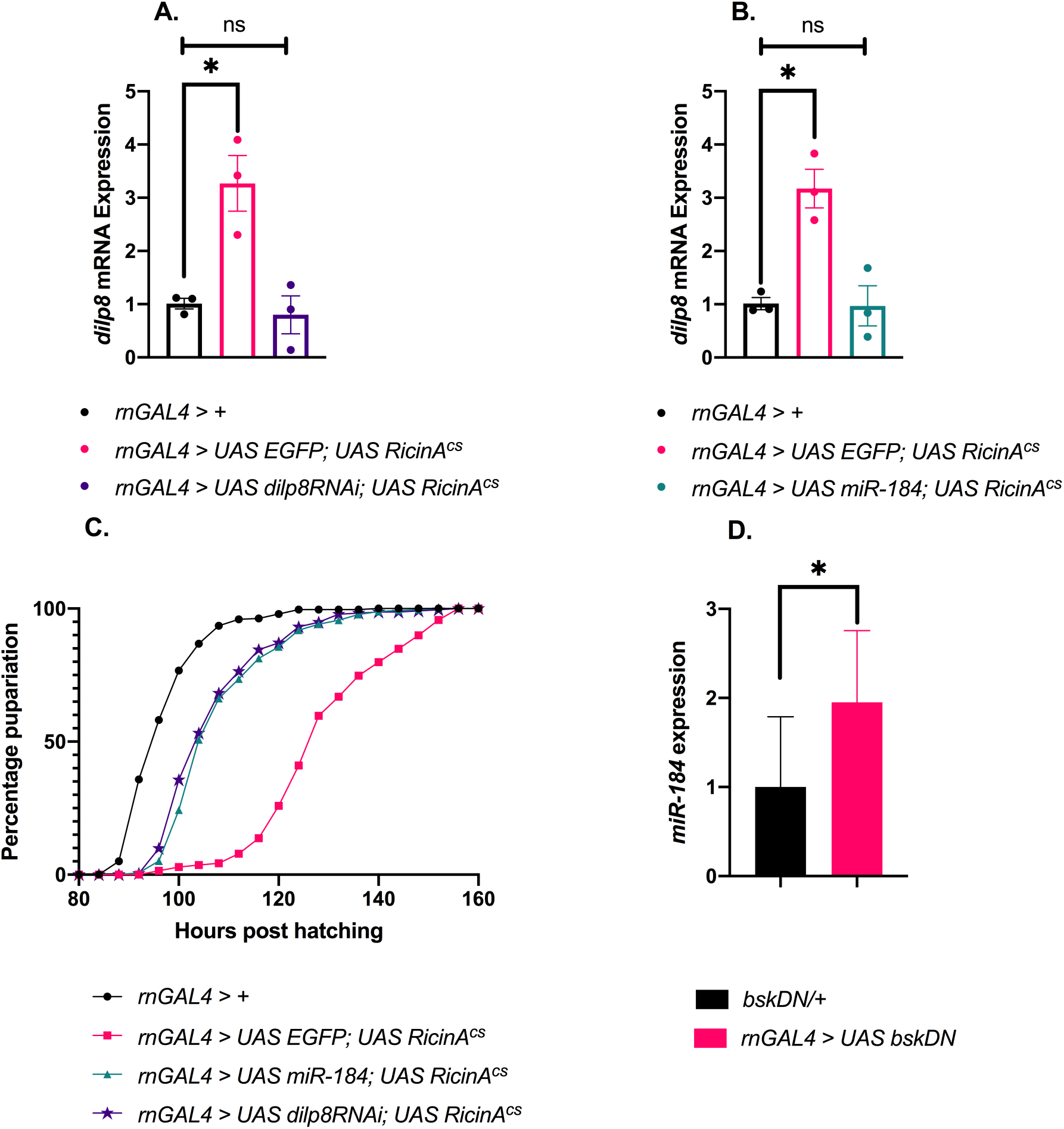
Role of miR-184 in Ricin-induced developmental perturbation. (A-B) Rescue of *dilp8* levels in the wing imaginal discs expressing Ricin-A, (A) *rnGal4* (control); *rnGal4, UAS-EGFP, UAS-RA^cs^* and *rnGal4, UAS-miR-dilp8-RNAi*, *UAS-RA^cs^,* three independent experiments are shown; (B) *rnGal4* (control); *rnGal4, UAS-EGFP, UAS-RA^cs^*; and *rnGal4, UAS-miR-184-OE, UAS-RA^cs^* three independent experiments are shown, For A and B, fold change of transcript levels of *dilp8* is shown, normalized to *rp49*, Total RNA was extracted from whole larvae. Bars graphs plotted are average/mean values, error bars represent SEM, a Brown-Forsythe and Welch’s ANOVA test was performed to determine statistical significance (C) Rescue of pupariation delays in response to Ricin-A expression in the imaginal discs, *rnGal4* (control) (n=298); *rnGal4, UAS-EGFP, UAS-RA^cs^* (n=139); *rnGal4, UAS-miR-184-OE, UAS-RA^cs^* (n=272) and *rnGal4, UAS-miR-dilp8-RNAi, UAS-RA^cs^*(n*=*233). Three independent experiments are shown; p<0.0001. Data points are percentage pupariated, LogRank test was performed to determine statistical significance. (D) Blocking JNK signaling in the imaginal discs leads to the induction of miR-184 levels, *UAS-bskDN* (control) and *rnGal4, UAS-bskDN,* four independent experiments are shown; p=0.0308. Fold change of transcript levels of *miR-184* is shown, normalized to *2SrRNA*. Total RNA was extracted from the wing imaginal disc. Bars graphs plotted are average/mean values. Error bars represent SEM. * p < 0.05, ***p<0.001, ****p<0.0001.

Finally, to investigate the signaling pathway that regulates *miR-184* expression under stress, we inhibited JNK signaling in the imaginal discs using the dominant-negative form of *basket*, *Drosophila* JNK (*bskDN*). We found that blocking JNK signaling increased *miR-184* levels in wing imaginal discs (Fig. 6D), confirming the role of JNK signaling in the regulation of *miR-184* levels and suggesting a similar response during tissue damage.

## Discussion

Achieving the final body size of an organism, as determined by its genetic makeup, requires precise coordination between growth and developmental timing. Environmental factors, which influence gene expression and signaling pathways that regulate these processes, introduce additional variability. To buffer against environmental fluctuations, biological systems employ robust mechanisms, including the post-transcriptional regulation of gene expression by microRNAs. A genetic screen conducted in our lab identified the microRNA, miR-184 as a regulator of pupariation timing. Among several *Drosophila* microRNAs, miR-184 is a maternally deposited microRNA (Lovino et al., 2009) with a highly dynamic expression pattern across various developmental stages (Li et al., 2011).

Here, we show that miR-184 is essential for normal ecdysone signaling and regulates the developmental transition from larval to pupal stages. The delayed pupariation observed in *miR-184* mutants was associated with reduced ecdysone signaling. Further, we show that *miR-184* acts in the imaginal discs to control ecdysone levels and pupariation timing. Our experiments identified *dilp8*, a gene encoding a paracrine factor that stabilizes organ and body size (Colombani et al., 2012; Garelli et al., 2012), as a direct target of *miR-184*. DILP8 delays pupariation by modulating ecdysone signaling during imaginal disc damage (Colombani et al., 2012; Garelli et al., 2012). Loss of *miR-184* resulted in elevated *dilp8* transcript levels, while suppressing *dilp8* in this context rescued the pupariation delay. Furthermore, we showed that miR-184 directly regulates *dilp8* expression via its 3’ untranslated region (UTR). Our findings <u>confirm</u> that *dilp8* levels are tightly controlled by miR-184 during normal larval development, with the loss of this regulation causing delays in pupariation.

Previous studies have highlighted the critical importance of maintaining optimal *dilp8* levels during development, as elevated *dilp8* expression is associated with pupariation delays. The role of DILP8 in ensuring developmental stability has been corroborated by several groups (Colombani, Andersen, & Leopold, 2012; Garelli et al., 2012; Jones et al., 2016). Notably, loss of *dilp8* increased wing asymmetry, underscoring its role in buffering developmental noise (Colombani, Andersen, & Leopold, 2012; Garelli et al., 2012, Garelli et al., 2015). Thus, we report a novel post-transcriptional mechanism by which *miR-184* regulates *dilp8* expression, a process essential for maintaining proper developmental timing.

The Hippo signaling pathway plays a pivotal role in maintaining *dilp8* expression during normal development, ensuring symmetric tissue growth (Boone et al., 2016). Additionally, during pupariation, ecdysone-triggered *dilp8* signaling from the cuticle epidermis orchestrates behavioral and morphogenetic transitions (Heredia et al., 2021). The post-transcriptional regulation of *dilp8* by Pacman/XRN1 (Jones et al., 2016) and the suppression of JNK signaling by dNOC1 (Destefanis et al., 2022) are also essential for maintaining appropriate *dilp8* levels during development. Building on this foundational work, our study identifies a novel post-transcriptional regulatory mechanism where *miR-184* precisely modulates *dilp8* expression, ensuring proper timing of critical developmental transitions.

Lgr3, the receptor for DILP8, is expressed in a small subset of neurons in the larval brain and plays a pivotal role in mediating developmental delays caused by DILP8 in response to tissue damage. The Lgr3 neurons directly interact with PTTH-producing neurons and insulin-producing neurons, forming a regulatory circuit that controls developmental timing and growth. Through the Lgr3-PTTH neuronal circuitry, DILP8 modulates ecdysone synthesis in the prothoracic gland, inducing developmental delays in response to tissue damage (Jaszczak et al., 2016). In this study, we show that the loss of *miR-184* elevates *dilp8* levels, which in turn reduces the expression of PTTH and disrupts ecdysone signaling. This cascade leads to the pupariation delays observed in *miR-184* mutants.

Recent studies have shown that in response to growth perturbations, DILP8 activates a developmental checkpoint that delays developmental timing and slows the growth of unaffected tissues to maintain overall organismal stability. Several signaling pathways regulate *dilp8* expression during tissue damage or stress. In early regenerating discs, JAK/STATsignaling induces *dilp8* expression, delaying the onset of pupariation in response to disc fragmentation (Katsuyama et al., 2015). Similarly, malignant disc tumors exhibit enhanced secretion of DILP8, regulated by JNK signaling (Romao et al., 2021). Blocking JNK signaling in damaged imaginal discs disrupts *dilp8* induction and alleviates eclosion delays (Colombani, Andersen, & Leopold, 2012; Katsuyama et al., 2015). In this study, we found that tissue damage leads to a reduction in *miR-184* levels, likely mediated by JNK activation, which in turn results in increased *dilp8* expression and developmental delays.

miR-184 plays a critical role in the female germline and early developmental processes (Lovino et al., 2009). miR-184 has also been identified as a nutrient-sensitive microRNA, with increased expression under low-nutrient conditions, where it is essential for survival (Fernandes et al., 2022; Gendron & Pletcher, 2017). Interestingly, overexpression of the epithelial tricellular junction protein Gliotactin upregulates *miR-184* levels via BMP signaling, establishing a feedback mechanism for its regulation (Sharifkhodaei et al., 2016). Furthermore, *miR-184* levels were observed to decrease in wing disc tumors (Shu et al., 2017), though the functional implications of this reduction remained unclear. These findings underscore the diverse regulatory roles of miR-184 in both normal and stress-induced developmental contexts.

*dilp8* levels in the imaginal discs of *Drosophila* serve as a critical developmental checkpoint (Colombani et al., 2012; Garelli et al., 2012). During regenerative growth, Dilp8, dMyc, and Wg levels are constrained by the *brain tumor* (*brat*) gene in regenerating tissues to ensure proper patterning, cell-fate specification, and differentiation (Abidi et al., 2023). Additionally, DILP8 plays a pivotal role during metamorphosis by maintaining the symmetry of adult organs (Boone et al., 2016; Garelli et al., 2012). Recent studies have demonstrated that DILP8, through its receptor Drl, is involved in the transdetermination of imaginal discs, highlighting its broader developmental significance (Nemoto et al., 2023). In adult *Drosophila*, *dilp8* levels in the ovarian follicle cells regulate ovulation (Liao, 2020). In *Aedes aegypti*, microRNA miR-277 has been shown to control lipid metabolism and reproduction by targeting *dilp8* (Ling et al., 2017). While our study establishes the role of *miR-184* in regulating *dilp8* in the context of imaginal disc damage, it remains an open question whether this regulatory axis extends to other developmental or physiological contexts. These findings suggest that *miR-184*-mediated control of *dilp8* may have broader implications in growth, regeneration, and tissue homeostasis across diverse systems.

Our study proposes a novel mechanism where *miR-184* fine-tunes *dilp8* levels to coordinate developmental timing by (i) maintaining appropriate ecdysone signaling under normal conditions and (ii) actively adjusting timing during perturbations to facilitate damage repair (Fig. 7A, B). These findings highlight the importance of post-transcriptional regulation in responding to developmental disturbances, opening avenues for future research on tissue-specific and cross-species mechanisms of microRNA-mediated regulation.

**Figure 7.**
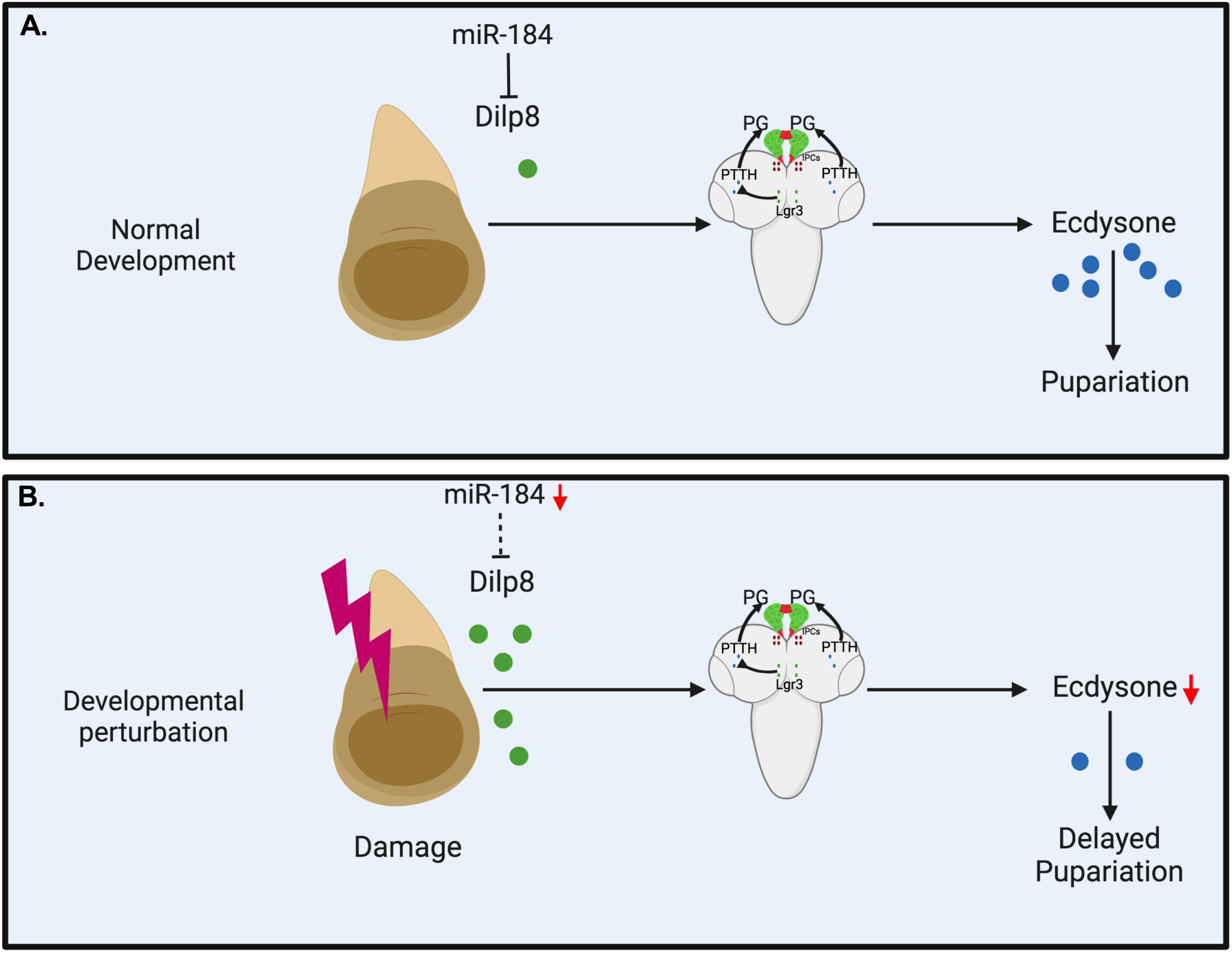
Model showing the function of miR-184 during growth and development. *miR-184* regulates *dilp8* expression to synchronize developmental timing by ensuring optimal ecdysone signaling under normal conditions (A) and by dynamically modifying this timing in response to perturbations to promote damage repair (B). Image created with BioRender.com.

### Experimental Methodology

#### Fly stocks

The fly stocks were reared in food containing standard laboratory cornmeal medium in an incubator maintained at 25°C, 70 % humidity, and a 12:12 Light: Dark (LD) cycle. 1 L of the *Drosophila* medium contains 58.2 g cornmeal, 50.8 g dextrose, 23.6 g yeast, 8 g agar, and 3 g nipagin (10% in 100% ethanol). *w^1118^* (6326), and *UAS-miR-184-sponge* (61396) lines were obtained from BDSC. The *miR-184^KO^* line (116326) was obtained from DGRC. The *UAS-Scramble-sponge* line was a kind gift from Dr. Tudor Fulga. The *UAS-dilp8RNAi* and *dilp8^MI00727^* lines were a kind gift from Dr. Pierre Leopold’s lab. The *tub-GAL4, phm-GAL4, aug21-GAL4, miR-10^KO^, miR-124^KO^, miR-133^KO^, miR-285^KO^, miR-283^KO^, miR-304^KO^, miR-210^KO^, miR-219^KO^, miR-31b^KO^, miR-137^KO^, miR-375^KO^, and miR-193^KO^* lines were a kind gift from Dr. Stephen Cohen’s lab. The *rn-GAL4* and *UAS-RA^cs^* lines were a kind gift from Dr. Marco Milan’s lab. The *ptth-GAL4* line was a kind gift from Dr. Nisha Kannan’s lab. The *UAS miR-184* line was a kind gift from Dr. Kweon Yu’s lab. The *dilp8 3’UTR-GFP* reporter line was created as described (Varghese & Cohen, 2007).

#### Pupariation time point assay

Pupariation time point assay measures the time taken from hatching to pupariation. All the larvae hatched before the peak hatching was picked and discarded. The remaining eggs are kept in a 25^°^C incubator for one hour. First instar larvae are collected within a tight time window of one hour. Collected larvae are reared in a 25^°^C incubator for the next few days, and the number of larvae pupariated is quantified every 4 hours. Logrank statistical Analyses were carried out in Oasis2 (Yang et al., 2016). Pupariation experiments involving the use of UAS-RA^cs^ were carried out at 29^°^C.

#### Ecdysone feeding experiment

L1 Larvae were reared in normal media at 25°C till 80 hrs post-hatching. At 80 hrs post-hatching, 25 larvae were transferred to either 20-hydroxyecdysone (0.3 mg/ml) (Sigma-Aldrich, H5142) or ethanol media and maintained at 25°C. The number of pupae was recorded every 4 hours.

#### Ecdysone quantification

For ecdysone extraction, 15 larvae were collected in 2 ml Safe-lock Eppendorf tubes, flash frozen in liquid nitrogen, and stored at -80°C at the indicated post-hatching timepoint. The samples were homogenized using 0.1mm Zirconium oxide beads and the Bullet Blender Storm (BBY24M from Next Advance) in 300 µl methanol (Sigma-Aldrich, 322415), centrifuged at 15000 rpm for 5 min at 4°C, following which the supernatant was transferred to a new microcentrifuge tube. 300 µl of methanol was added to the samples and vortex mixed before centrifuging and transferring the supernatant to the same tube. Lastly, the procedure was repeated with 300 µl of ethanol (Himedia, MB228), to obtain a final volume of 900 µl of the supernatant. The samples containing the supernatant were centrifuged at 15000 rpm for 5 min at 4°C in multiple rounds while transferring the supernatant to remove any remaining debris. The samples were evaporated using an Eppendorf Speedvac centrifuge and the pellet was dissolved in 200µl of ELISA buffer (EIA Buffer). The samples were stored at room temperature for 2 hours with intermediate vigorous vortexing to ensure proper dissolution of the pellet. ELISA was performed as per the instructions of the 20-Hydroxyecdysone ELISA kit (Cayman Chemicals, 501390). The readings were taken at 405 nm using the TECAN Infinite M200 pro-multimode plate reader. The data was analysed as per the manufacturer’s instructions.

#### Sample collection for RNA isolation

Larval samples were collected 116 hrs post-hatching and plunged into liquid nitrogen. The samples were then transferred to -80^°^C for storage. For the imaginal discs RNA isolation: 20 Larvae per set were collected ∼92-100 hrs post-hatching and dissected in PBS solution. The wing imaginal discs were isolated and stored in TriZol immediately post-dissection before proceeding for total RNA extraction.

#### qRT-PCR

Total RNA was extracted from samples each using the TriZol Reagent, as per the manufacturer’s instruction, and pelleted using chloroform and isopropanol. The pellet was washed with 70% ethanol and centrifuged at 15,000 rpm. The pellet was dried and dissolved in 40µl of Molecular biology-grade water (Himedia). Any contaminating DNA was degraded with the help of a QIAGEN RNase-Free DNase set as per the manufacturer’s instructions. Following this, the RNA was re-extracted and dissolved in 40µl of Molecular biology-grade water (Himedia). Takara Perfect Real-time kit was used to convert similar quantities of the RNA to cDNA. The cDNA was used for qPCR quantification using TB Green Premix Ex Taq (Tli RNaseH Plus) in a CFX96 machine (Bio-Rad). The level of the genes was normalized to the level of actin. The .LCt values generated were analysed using the Student’s t-test in GraphPad Prism.

#### microRNA qRT-PCR

Total RNA was extracted from L3 larvae using the TriZol extraction method (described above). miR-184 was reverse transcribed and the qPCR was performed as per instructions of the Takara miR-X kit.

Primers used for qRT-PCRs

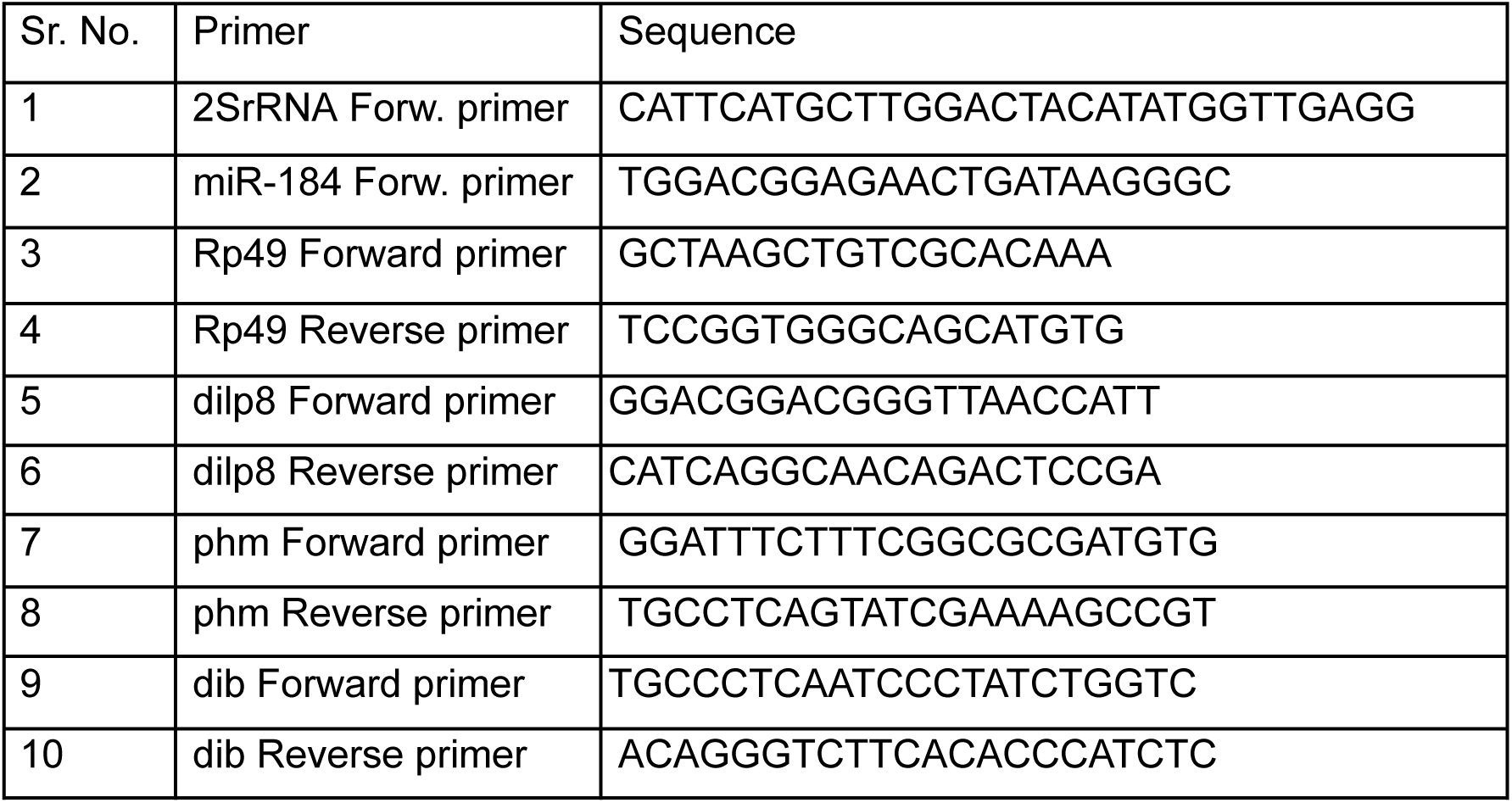
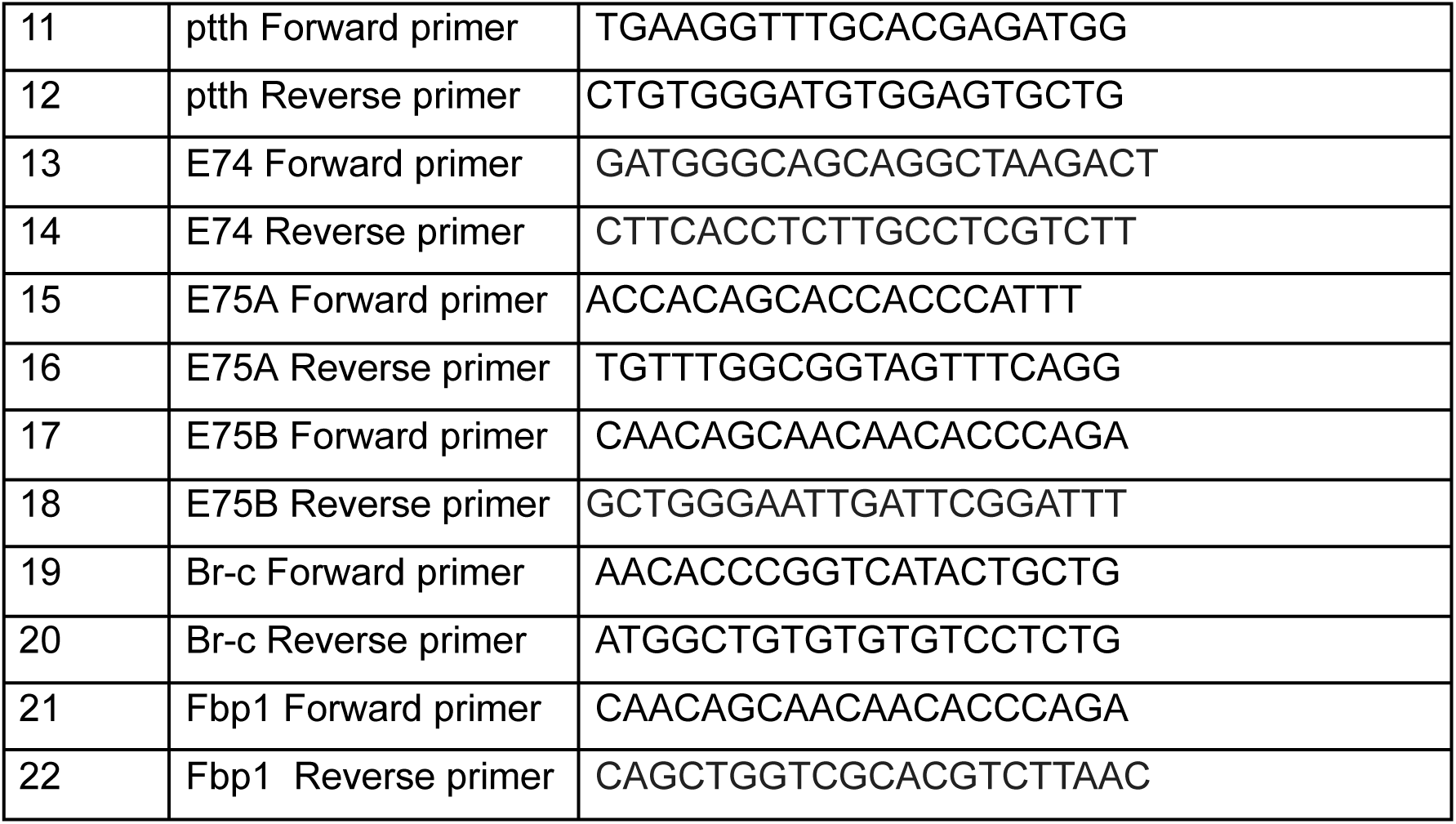

### Antibody stainings and immunofluorescence

For every genotype, non-wandering early third instar larvae were dissected in 1x ice-cold PBS and fixed in 4% paraformaldehyde (PFA) (Cat# P6148, Sigma-Aldrich) solution for 20 minutes at RT. Following the removal of PFA, the tissues were washed using PBT (1x phosphate-buffered saline + 0.1% Triton X-100 [Bio-Rad Cat. No. 161-0407]). After adding the blocking solution (PBT + 0.1% BSA [Cat. no. A2153 from Sigma-Aldrich]), the samples were allowed to sit at room temperature for forty-five minutes on a nutator to ensure efficient blocking. The samples were then incubated with anti-ptc antibody (DSHB antibody 1:200 dilution with blocking solution) and anti-GFP antibody (polyclonal antibody 1:500 dilution with blocking solution) (Cat. no. A-6455 from Invitrogen) overnight at 4°C while being rotated continuously. Following a thorough PBT wash, the tissues were left to incubate for two hours at room temperature with the secondary antibody. Anti-mouse IgG-conjugated with Alex-Fluor 633 and anti-rabbit IgG-conjugated with Alex-Fluor 488 diluted with blocking solution were used to fluorescently label the samples respectively. The samples were thoroughly cleaned and mounted using a drop of Thermo Fisher Scientific’s SlowFade Gold Antifade Reagent (Cat. no. S36939) after two hours. The samples were visualized and imaged by a confocal microscope (Leica DM6000B). ImageJ software was used to analyse the images further.

### Bioinformatics

microRNA targets were predicted using the target prediction software, TargetScan Fly 7.2 and DIANA-microT-CDS. STarMir was further used to confirm and generate the RNA hybrid (Rennie et al., 2014).

## ACKNOWLEDGMENTS

We are thankful to Dr. Smitha Vishnu for her critical comments. This work was supported by intramural funds from IISER Thiruvananthapuram. This work was also supported by a Ramanujan Fellowship to JV from Science and Engineering Research Board (SERB), Department of Science and Technology (DST) - (SR/S2/RJN-140/2011); Extra Mural Research Grant from Science and Engineering Research Board (SERB), Department of Science and Technology (DST) - (EMR/2016/004978); Core Research Grant, Science and Engineering Research Board (SERB), Department of Science and Technology (DST) - (CRG/2023/002329); and Research Grant under Scheme for Transformational and Advances Research in Sciences (STARS), by Indian Institute of Science (IISc), Ministry of Education (MoE) (MoE-STARS/STARS-2/2023-0108). We also thank The University Grants Commission (UGC), India for providing Junior and Senior Research Fellowship to JF and the Council for Scientific and Industrial Research (CSIR), India for providing a Junior Research Fellowship for MN.

## AUTHOR CONTRIBUTIONS

Conceptualization: JF and JV; Methodology: JF, MN, AMMP and JV; Investigation: JF, MN and AMMP; Data acquisition: JF, MN and AMMP; Data analysis: JF and MN; Writing - Original Draft: JF; Writing - Review & Editing: JV; Funding Acquisition: JV; Supervision: JV.

