## Supplemental legend for "miR-184 modulates *dilp8* to control developmental timing during normal growth conditions and in response to developmental perturbations"

### Supplementary legends:

**Table S1: Genetic screen to identify microRNAs that play a role in regulating pupariation timepoint**

Mean pupariaition time for each genotype is plotted. Total number of larvae > 100 except for the *miR-133*$^{KO}$ (n=70) and *miR-193*$^{KO}$ (n=85) genotypes. p<0.0001 for all genotypes. Log-rank test was performed to determine statistical significance.

**Figure S1. *miR-184*-sponge efficiency**

*miR-184 levels* when *miR-184* was downregulated in the whole larvae using *tubGAL4*; *tubGAL4 > UAS Scramble* (control) and *tubGAL4 > UAS miR-184-sp*. Data from three independent experiments are shown; p=0.0386. Fold change of *miR-184* is shown, normalized to *2SrRNA*. Total RNA was extracted from the whole larvae. Bars graphs plotted are average/mean values, error bars represent SEM, and a Welch's t-test was carried out to determine statistical significance

**Figure S2. *miR-184*-sponge specificity**

Rescue of pupariation delay, *tubGal4, UAS Scramble* (control) (n=184) and *tubGal4, UAS miR-184-sp* (*miR-184* downregulation) (n=59) and *tubGal4, UAS miR-184OE, UAS miR-184 sp* (n=188), three independent experiments are shown. Data points plotted are percentage pupariated, LogRank test was performed to determine statistical significance.

**Figure S3. *miR-184* levels in the imaginal discs maintain proper Ecdysone signaling**

(A) Expression of Ecdysone biosynthesis genes in response to imaginal disc downregulation of *miR-184*; using *rnGal4; UAS miR-184-sp* and *rnGal4, UAS Scramble* (control), from four independent experiments are shown. Fold change of transcript levels of *ptth*, *phm*, and *dib* are shown, normalized to *rp49*. Total RNA was extracted from whole larvae.

(B) Expression of Ecdysone response genes in response to imaginal disc downregulation of *miR-184* using *rnGal4*; *rnGal4, UAS Scramble* (control) and *rnGal4, UAS miR-184-sp,* data from four independent experiments are shown. Fold change of transcript levels of *E74*, *E75A*, *E75B*, *BR-C*, and *fbp1* are shown, normalized to *rp49*. Total RNA was extracted from whole larvae. Bars graphs plotted are average/mean values, error bars represent SEM; a two-way ANOVA was performed to determine statistical significance; * p < 0.05, ***p<0.001, ****p<0.0001.

**Figure S4. *miR-184* target site in the *dilp8-3'UTR***

Image generated by STarMir showing the binding between miR-184 mature sequence and the target sequence in the *dilp8 3'UTR*. Seed sequence is shown in red. Minimum free-energy value of the hybridization is also shown.
