## Supplementary material for "miR-184 modulates *dilp8* to control developmental timing during normal growth conditions and in response to developmental perturbations": Table S1

| Table S1 | |
|---|---|
| **microRNA (KO/KO)** | **Mean Pupariation time (hph)** |
| miR-10 | 128 |
| miR-124 | 132 |
| miR-133 | 196 |
| miR-285 | 132 |
| miR-283 | 136 |
| miR-304 | 124 |
| miR-210 | 128 |
| miR-219 | 124 |
| miR-31b | 140 |
| miR-137 | 148 |
| miR-375 | 120 |
| miR-193 | 132 |
| miR-184 | 132 |
| w1118 | 118 |
