## Supplementary figures and images for "miR-184 modulates *dilp8* to control developmental timing during normal growth conditions and in response to developmental perturbations"

### Figure S1

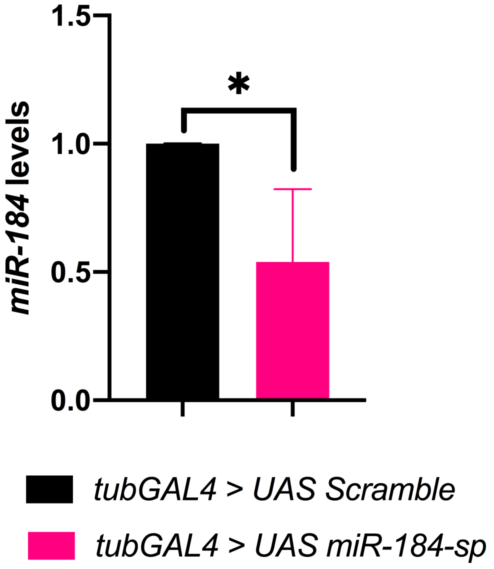

### Figure S2

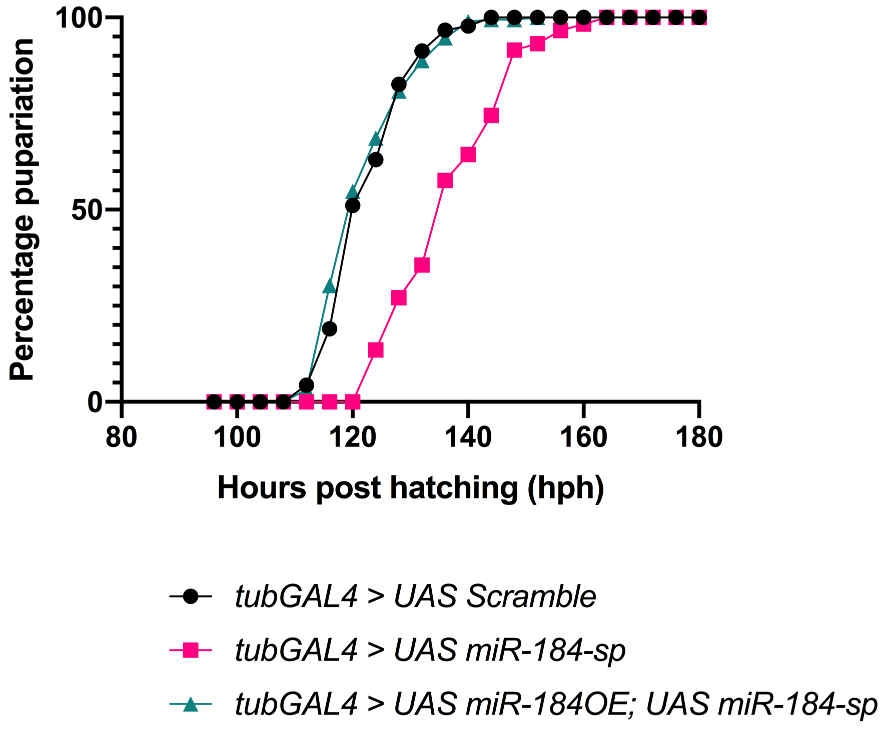

### Figure S3

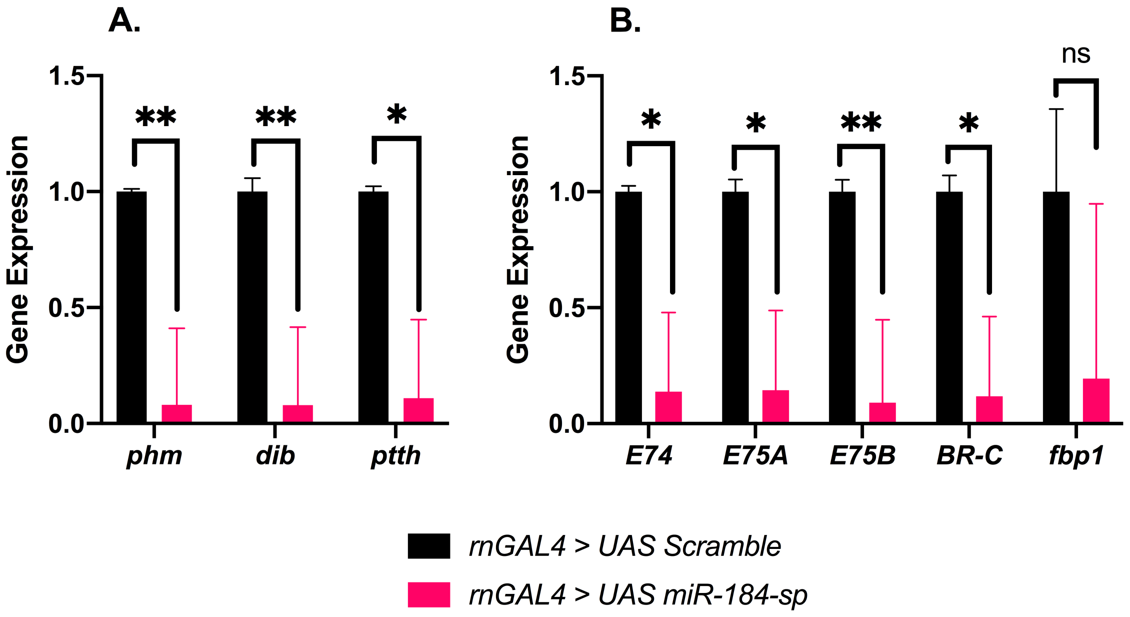

### Figure S4

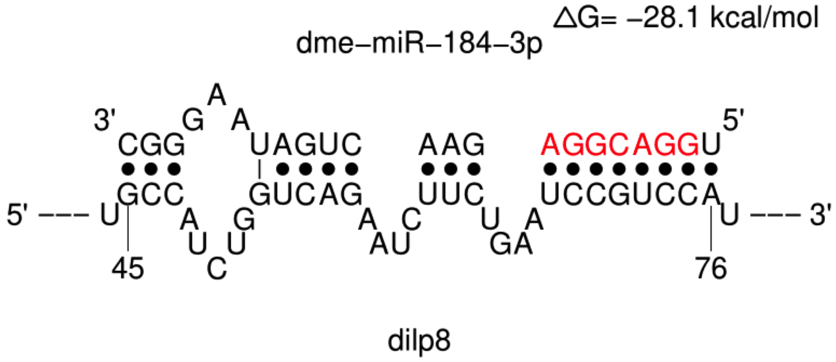
